# Women at the receiving end: Exploring Couples’ experiences of infertility challenges in Nigeria

**DOI:** 10.1101/2025.01.08.632060

**Authors:** Deborah Tolulope Esan, Kelechukwu Queendaline Nnamani, Adetunmise Oluseyi Olajide, Oluwadamilare Akingbade, David Bamidele Oluwade, David Bamidele Olawade, Carlos Guillermo Ramos

## Abstract

**Background:** Infertility remains a significant global health issue with profound emotional and social consequences, particularly in patriarchal societies. This study explores the lived experiences of infertile couples in Nigeria, focusing on societal perceptions, gendered blame, and the role of spirituality.

**Methods:** A qualitative exploratory design was adopted, with data collected from purposively sampled infertile couples attending the Federal Teaching Hospital, Ido-Ekiti, Nigeria. Semi-structured interviews were conducted, and thematic analysis was used to identify recurring patterns and themes.

**Results:** Thematic analysis identified two main themes and seven sub-themes. The first theme, Couples**’** Perception of Infertility, included five sub-themes: definition of infertility, perceived causes, perceived types, infertility as a feminine issue, and the role of spirituality. The second theme, Challenges of Couples Experiencing Infertility, comprised two sub-themes: women at the receiving end and poor societal support.

**Conclusion:** The findings shows the interplay of medical, cultural, and spiritual dimensions in shaping infertility perceptions. The study highlights the disproportionate burden on women and inadequate societal support, calling for public health interventions to address stigma and promote gender-inclusive reproductive health education.

## Introduction

Infertility, defined as the inability to conceive after one year of regular unprotected intercourse, affects approximately 15% of couples globally, making it a significant public health challenge (1,2). In many societies, infertility is not just a biological condition but a deeply personal, cultural, and societal issue, with far-reaching implications for the emotional and social well-being of affected individuals. This is especially true in Nigeria, where cultural norms and expectations place a high premium on childbearing, often resulting in women bearing a disproportionate share of blame and stigma associated with infertility (3,4).

Infertility is perceived through a mix of medical, spiritual, and societal lenses. Medically, causes such as hormonal imbalances, infections, or fibroids are widely recognized (5). However, cultural beliefs and spiritual explanations, such as witchcraft, curses, or divine punishment, often shape couples’ understanding of infertility, creating a dual explanatory framework that intertwines the biomedical with the supernatural. This combination influences how couples interpret their experiences and navigate societal expectations (6,1).

Despite the acknowledgment that both men and women can contribute to infertility, Nigerian society overwhelmingly perceives it as a feminine issue, further intensifying the gendered burden. Women are often blamed, ostracized, or even abandoned due to their perceived inability to fulfill societal and familial expectations of motherhood (7,4). This blame persists even when male factors, such as low sperm count, are identified, reflecting deeply ingrained gender norms. As participants in this study noted, infertility-related stigma often leads to emotional and social isolation for women, exacerbated by limited societal support.

The consequences of infertility extend beyond personal suffering to impact couples’ relationships and community standing. Emotional distress, marital strain, and social ridicule are common experiences for couples navigating infertility (8, 5). Women, in particular, face societal pressures that amplify their challenges, including ridicule, unsolicited advice, and even suggestions of seeking children outside their marriages (4). These societal attitudes not only diminish the well-being of affected individuals but also highlight the inadequacies of existing support systems.

The interplay of medical, cultural, and spiritual perspectives creates a complex landscape in which couples must navigate their infertility experiences. Previous research highlights that inadequate societal support and a lack of understanding exacerbate the challenges faced by affected couples, leaving them vulnerable to stigma and emotional distress (6,2). This study delves into these issues, exploring the gendered dimensions of infertility, societal perceptions, and the pressing need for supportive interventions in the Nigerian context.

### Aim of the Study

This study aims to examine the lived experiences of couples dealing with infertility, focusing on the societal and cultural factors that shape their journeys. Specifically, it highlights the disproportionate burden borne by women and the inadequacy of societal support systems. By shedding light on these issues, the study seeks to foster a deeper understanding of infertility as a multidimensional challenge and advocate for more inclusive and empathetic societal responses.

## Materials and Methods

### Research Design

This study employed a qualitative exploratory research design to explore the complexities and nuanced experiences surrounding infertility among couples. This approach facilitated in-depth conversations with participants, enabling a comprehensive understanding of their perceptions and challenges.

### Study Location

The study was conducted at the Federal Teaching Hospital, Ido-Ekiti, located in the Ido-Osi Local Government Area of Ekiti State, Nigeria. This 280-bed tertiary healthcare institution, formerly known as the Federal Medical Centre, serves as a referral center for the region.

### Study Population

The target population comprised couples who had been attempting to conceive for at least 12 months without success and were receiving treatment at the infertility clinic of the hospital’s Obstetrics and Gynecology unit.

### Sampling Method

Purposive sampling was used to recruit participants with rich and relevant insights into the research phenomenon. Inclusion criteria were:

1. Couples with primary or secondary infertility diagnosed after at least 12 months of trying to conceive.
2. Couples actively undergoing treatment at the hospital and willing to participate during the study period.

### Data Collection Procedure

The data collection was between January 2019 and April 2019. Face-to-face, semi-structured interviews were conducted during clinic visits. After explaining the study’s purpose and obtaining informed consent, interviews were conducted in private clinic spaces to ensure confidentiality. Each session lasted 30-40 minutes, using an open-ended interview guide to foster flexibility. Interviews were recorded, transcribed verbatim, and analyzed manually.

### Data Analysis

Thematic analysis was employed in six phases: familiarization, coding, theme generation, theme review, theme definition, and write-up. Open coding was used to conceptualize and categorize data. Quantitative data were analyzed to describe participants’ demographic profiles.

### Ethical Approval and Consent

Ethical approval was obtained from the Ethics and Research Committee of the Federal Teaching Hospital, Ido-Ekiti (Protocol Number: ERC/2018/11/26/158B). Participants in this study provided verbal informed consent, which was documented in written notes by the researchers, with a witness present to ensure ethical compliance and accuracy. Participants’ rights and confidentiality was ensured throughout the study.

### Results

The table 1 shows that most participants are aged 20-35, with all identifying as Christian. The Yoruba ethnicity and married status dominate the group. Employment rates are high, although a significant proportion have no children, indicating the context of infertility among participants

**Table 1:** Summary of Socio-Demographic Profile of Participants.

| Socio-Demographic Data | Frequency (n=15) | Percentages (%) |
| --- | --- | --- |
| Age in years |  |  |
| 20-35 | 12 | 80.0 |
| 36 and above | 3 | 20.0 |
| **Religion** |  |  |
| Christianity | 15 | 100.0 |
| Islam | - | - |
| Others | - | - |
| **Ethnicity** |  |  |
| Yoruba | 12 | 80.0 |
| Hausa | - | - |
| Igbo | 1 | 6.7 |
| Others | 2 | 13.3 |
| **Marital Status** |  |  |
| Married | 14 | 93.3 |
| Separated | 1 | 6.7 |
| Divorced | - | - |
| Widow | - | - |
| **Employment Status** |  |  |
| Employed | 13 | 86.7 |
| Unemployed | 2 | 13.3 |
| **Years of Marriage** |  |  |
| 1-5 | 8 | 53.3 |
| 6-10 | 5 | 33.3 |
| 11 and above | 1 | 6.7 |
| **Number of Children** |  |  |
| Nil | 10 | 66.7 |
| 1-2 | 5 | 33.3 |
| 3 and above | - | - |
| **Sex** |  |  |
| Male | 2 | 13.3 |
| Female | 13 | 86.7 |

**Table 2:** Themes and sub-themes generated.

| Serial Number | Themes | Sub-Themes |
| --- | --- | --- |
| 1 | Couples' Perception of Infertility |  |
| 1.1 |  | Definition of infertility |
| 1.2 |  | Perceived causes of infertility |
| 1.3 |  | Perceived types of infertility |
| 1.4 |  | Infertility as a feminine issue |
| 1.5 |  | The role of spirituality in infertility |
| 2 | Challenges of Couples Experiencing Infertility |  |
| 2.1 |  | Women at the receiving end |
| 2.2 |  | Poor societal support |

### Theme 1: Couples’ Perception of Infertility

Infertility is a deeply personal and culturally influenced phenomenon, often defined and interpreted through a mix of medical, spiritual, and societal lenses. Couples in this study expressed varied perspectives about what infertility entails, its causes, and its implications. These perceptions are critical in shaping how couples understand their condition and engage with treatments and societal expectations. This theme explores the definitions, perceived causes, and types of infertility, as well as the gendered blame and the role of spirituality in shaping beliefs.

#### Sub-Themes

##### 1. Definition of Infertility

Participants largely defined infertility as the inability to conceive, highlighting the challenges faced in achieving natural conception. For some, infertility was explained in layman’s terms, while others demonstrated an understanding of its medical dimensions. This understanding formed the foundation for their interpretations and coping mechanisms. This is supported by the following quotes:

> *“Infertility is the inability to conceive.” (Participant 1, female, 30 years old, married for 2 years, has no child)*.
>
> *“Infertility is when a woman is unable to conceive; that is, when someone is unable to get pregnant naturally, or through sexual intercourse.” (Participant 2, female, 35 years old, married for 8 years, has no child)*.
>
> *“Infertility is when you are unable to conceive; maybe the fault is from the man or the woman, and many things can cause that—maybe an infection, fibroid, or for the man, low sperm count.” (Participant 6, female, 35 years old, separated, has a child)*.

##### 2. Perceived Causes of Infertility

Participants identified multiple causes of infertility, encompassing both medical and spiritual explanations. Medical causes included infections, fibroids, hormonal imbalances, and maternal age. However, several participants attributed infertility to supernatural causes, such as witchcraft, curses, or divine punishment. This dual attribution reflects a blend of cultural beliefs and biomedical knowledge. This perspective is captured in these direct quotes

> *“I think infection is a cause; also, some say age.” (Participant 1, female, 30 years old, married for 2 years, has no child)*.
>
> *“Hormonal imbalance could cause it. Some even say it runs in families. Some people spend years before having a child, so I think it can be a family issue.” (Participant 2, female, 35 years old, married for 8 years, has no child)*.
>
> *“In my case, they said fibroid. Fibroid can cause infertility and some other infections.” (Participant 4, female, 38 years old, married for 5 years, has no child)*.
>
> *“I don’t blame anybody. I count it as part of the journey of life, a challenge as a human being.” (Participant 2, female, 35 years old, married for 8 years, has no child)*.
>
> *“I did one abortion; that’s why.” (Participant 7, female, married for 7 years, has a 15-year-old child)*.
>
> *“There are evil people in this world who don’t want people to marry or have children.” (Participant 1, female, 30 years old, married for 2 years, has no child)*.
>
> *“Ah yes, nah. With experience, witches can cause infertility. I have seen someone say, ‘You won’t have a child because you married my husband.’” (Participant 5, female, 48 years old, married for 8 years, has a 5-year-old child)*.

##### 3. Perceived Types of Infertility

Participants demonstrated an understanding of the different types of infertility, describing primary infertility as the inability to conceive at all, secondary infertility as difficulty with subsequent pregnancies, and sub-infertility or pregnancy wastage as recurrent miscarriages.

This remarks from participant further exemplify this:

> “*Primary infertility is when the couple or the woman has not given birth before. Secondary infertility is when there is an issue with subsequent pregnancies. Sub-infertility or pregnancy wastage is when pregnancies are lost through miscarriages.”* (Participant 8, female, 30 years old, married for 2 years, has no child).

##### 4. Infertility as a Feminine Issue

In Nigerian society, infertility is overwhelmingly perceived as a problem that affects women, despite the recognition that men can also contribute to infertility. This gendered blame leads to social and emotional burdens for women, even when male factors are involved. This point is highlighted by these participant responses:

> “*This part of the world, the first person to blame is the woman because of our belief.” (Participant 8, female, 30 years old, married for 2 years, has no child)*.
>
> *“Most times, it’s the woman.” (Participant 10, female, 36 years old, married for 10 years, has a 10-year-old child)*.
>
> *“In Yoruba culture, everything concerning infertility is blamed on the woman. They say maybe it’s due to many abortions or spiritual causes.” (Participant 2, female, 35 years old, married for 8 years, has no child)*.

##### 5. The Role of Spirituality in Infertility

Spiritual beliefs and practices significantly influenced participants’ perceptions of infertility. While some participants relied on medical explanations, others believed strongly in spiritual causes, such as witches, wizards, or divine punishment. This is echoed in the following participant statements

> “*Sometimes it may be spiritual, and sometimes it may not be.” (Participant 2, female, 35 years old, married for 8 years, has no child)*.
>
> *“There are evil people who don’t want others to have children.” (Participant 5, female, 48 years old, married for 8 years, has a 5-year-old child)*.

### Theme 2: Challenges of Couples Experiencing Infertility

Infertility poses numerous challenges for couples, with women bearing a disproportionate share of the blame and stigma. This is compounded by inadequate societal support systems, leaving many couples isolated and vulnerable. The theme explores the emotional, social, and cultural difficulties faced by individuals and highlights the pressing need for supportive interventions.

#### Sub-Themes

##### 1. Women at the Receiving End

Women were consistently identified as the primary recipients of blame for infertility. Participants noted that cultural and familial pressures often placed women at the center of criticism, even when male factors were involved. This gendered burden exacerbated the emotional and social challenges faced by women. This perspective is captured in these direct quotes.

> *“Most times, it’s the woman that is blamed for infertility. She is seen as the one having problem” (Participant 10, female, 36 years old, married for 10 years, has a 10-year-old child)*.
>
> *“In Yoruba culture, all the blame is on the woman. They say maybe it’s due to many abortions or spiritual causes.” (Participant 2, female, 35 years old, married for 8 years, has no child)*.
>
> *“You see some women being dragged out of the home because of this issue.” (Participant 5, female, 48 years old, married for 8 years, has a 5-year-old child)*.

##### 2. Poor Societal Support

Participants described a lack of societal understanding and support for couples experiencing infertility. Societal attitudes often amplified the stress and isolation felt by couples, with women facing ridicule and unhelpful advice, such as seeking children outside their marriages. Participant narratives support this, as seen in the following quotes:

> *“They are not getting any support at all; people just advise to impregnate someone outside.” (Participant 6, female, 35 years old, separated, has a 16-year-old child)*.
>
> *“People are not supportive at all; they just talk anyhow about you.” (Participant 10, female, 36 years old, married for 10 years, has a 10-year-old child)*.

## Discussion

The socio-demographic profile of participants in this study provides valuable insights into the context of infertility in a Nigerian setting. The predominance of female participants (86.7%) reflects the societal framing of infertility as a primarily feminine issue, consistent with cultural norms that disproportionately place the burden of childbearing on women (2,3, 7). This gendered narrative highlights the need for inclusive approaches in addressing infertility, emphasizing both male and female contributions to reproductive challenges (1, 9).

The age distribution, with 80% of participants aged 20–35, suggests that infertility is most commonly addressed during peak reproductive years. This aligns with global patterns where individuals within this age range actively seek medical or alternative solutions to fertility issues (8,9). However, the presence of older participants (20%) shows the persistent stigma surrounding infertility, compelling couples to continue seeking solutions despite advancing age (6).

The religious homogeneity, with all participants identifying as Christian, and the dominance of the Yoruba ethnic group (80%), reflect the study’s geographic location and cultural setting. These factors are critical in understanding how religious and cultural beliefs shape perceptions of infertility. For example, spiritual explanations such as witchcraft or divine punishment often coexist with biomedical views, influencing how individuals approach treatment and support-seeking behaviors (1,10).

The marital status data reveal that most participants are married (93.3%), with only one participant reporting separation. This is expected, as infertility is primarily a concern within marital contexts where societal and familial expectations for childbearing are most pronounced (11, 12). The single separated participant highlights the potential marital strain and breakdown that infertility can cause, a phenomenon frequently reported in patriarchal societies.

Employment data show that most participants are employed (86.7%), which could indicate a certain level of socio-economic stability among the study group. However, the emotional and financial burden of infertility treatments, compounded by societal stigma, remains a challenge (7). The 13.3% unemployment rate, particularly among female participants, may reflect broader gender disparities in employment opportunities, exacerbating the vulnerability of women facing infertility The majority of participants (66.7%) reported no children, consistent with the inclusion criteria focused on infertility. However, the 33.3% with one or two children highlight secondary infertility, which can carry an equally heavy emotional toll (13). The number of years of marriage also varied, with over half (53.3%) married for 1–5 years. This group likely represents couples in their early efforts to conceive, contrasting with those married for longer periods, who may face compounded emotional and social pressures (6).

The socio-demographic profile of participants reveals how cultural, gendered, and socio-economic factors intersect to shape the lived experiences of infertility. These findings emphasize the importance of context-specific interventions that address the unique challenges faced by individuals and couples within this setting. Public health initiatives should aim to mitigate stigma, foster equitable support systems, and promote inclusive reproductive health education that integrates cultural sensitivities (4,9).

This study highlights the complex and multidimensional nature of infertility, encompassing biomedical, spiritual, and societal dimensions. Participants’ definitions of infertility ranged from basic understandings as the inability to conceive to more nuanced interpretations incorporating cultural and medical insights. This variability shows both awareness and knowledge gaps regarding infertility, particularly its complexities. These findings align with previous studies, such as one in Ethiopia, which reported that participants had a general understanding of infertility but lacked clarity about specific contributing factors (8). Similarly, in India, participants recognized infertility as a medical condition but demonstrated limited understanding of its biological underpinnings (9).

Participants also articulated a dual explanatory model of infertility, attributing its causes to both medical and spiritual factors. Medical explanations included infections, fibroids, hormonal imbalances, and maternal age, while spiritual causes encompassed witchcraft, curses, and divine punishment. This duality mirrors findings from Uganda, where infertility was attributed to both biomedical and supernatural factors (10). In rural China, similar spiritual beliefs led individuals to combine traditional healing with medical treatments (11). These parallels emphasize the global prevalence of culturally specific narratives that coexist with biomedical explanations, reflecting the sociocultural context in which infertility is understood and addressed.

Participants demonstrated an awareness of infertility’s types—primary infertility (no prior pregnancies), secondary infertility (difficulty conceiving after a previous pregnancy), and sub-infertility (recurrent miscarriages). This nuanced understanding is not in tandem with findings from Japan and Nigeria where less than average of the participants has awareness of infertility (14, 15).

Despite recognizing male contributions to infertility, participants overwhelmingly reported that societal blame disproportionately falls on women. This gendered narrative is entrenched in patriarchal norms, which hold women solely responsible for childbearing. This mirrors findings from United state and in most part of the world where infertility was framed as a woman’s issue, with men rarely held accountable (16-18).

Spiritual interpretations of infertility also emerged as a prominent theme. Participants linked infertility to witchcraft, curses, or divine punishment, reflecting the cultural integration of spirituality and reproductive health in Nigeria. These beliefs are consistent with studies that identified spiritual explanations for infertility, attributing it to divine punishment, witchcraft, or the dissatisfaction of ancestors (19-24). These findings highlight the enduring influence of cultural and spiritual frameworks on health-seeking behaviors.

Women’s experiences were marked by significant emotional and social burdens due to infertility. Participants noted that women faced blame, ostracism, and in some cases, domestic violence. These findings resonate with studies from Ghana, Zimbabwe and Nigeria, where societal norms perpetuated the stigmatization of infertile women (25-27). A pervasive lack of societal support compounded the challenges faced by couples. Participants described experiencing ridicule and unhelpful advice, which heightened their emotional stress. Research has shown that strong societal support significantly enhances the coping ability of individuals experiencing infertility (28, 29).

### Implications for Practice and Research

These findings shows the need for comprehensive public health interventions that address both the biomedical and socio-cultural dimensions of infertility. Health education campaigns should focus on dispelling myths surrounding infertility while promoting awareness of male factors and reducing the stigma faced by women. Policies that integrate community-based support systems, including counseling and peer support groups, could mitigate the social and emotional challenges encountered by affected couples.

Future research should explore the intersection of cultural, spiritual, and medical narratives in shaping infertility experiences across diverse contexts. Longitudinal studies could provide deeper insights into how these narratives influence coping mechanisms and treatment-seeking behaviors over time. Expanding this research to include male perspectives could also address the gender imbalance in current narratives and promote a more equitable understanding of infertility.

By contextualizing infertility as a multidimensional issue, this study contributes to a growing body of literature that advocates for culturally sensitive and inclusive approaches to reproductive health.

## Conclusion

This study shows the intricate and multidimensional challenges associated with infertility, as shaped by a confluence of biomedical, cultural, and spiritual dimensions. Participants’ narratives revealed significant awareness of infertility’s causes and types but also highlighted critical knowledge gaps and the persistent influence of spiritual and cultural beliefs. A dual explanatory model blending medical and spiritual frameworks was prevalent, reflecting how couples navigate their infertility journeys within sociocultural contexts.

Gendered stigma emerged as a dominant theme, with women bearing disproportionate blame and suffering due to entrenched patriarchal norms. The societal perceptions of infertility as a feminine issue exacerbate emotional distress and social isolation, particularly for women, despite acknowledgment of male infertility contributions. Additionally, the lack of robust societal support systems perpetuates stigma and psychological challenges for affected couples.

Addressing these issues requires multifaceted interventions that incorporate culturally sensitive health education, equitable access to diagnostic and treatment services, and supportive community frameworks. By fostering understanding and inclusivity, these efforts can mitigate the societal burden of infertility, promote gender equity, and enhance the well-being of affected individuals and couples.

## Acknowledgements

We sincerely thank all participants for their invaluable contributions and for courageously sharing their experiences despite the sensitive nature of infertility. Their openness has significantly enriched our understanding of this critical issue in Nigeria.

